# SpiDa-MRI, behavioral and (f)MRI data of adults with fear of spiders

**DOI:** 10.1101/2024.02.07.578564

**Authors:** Mengfan Zhang, Alexander Karner, Kathrin Kostorz, Sophia Shea, David Steyrl, Filip Melinscak, Ronald Sladky, Cindy Sumaly Lor, Frank Scharnowski

## Abstract

Neuroimaging has greatly improved our understanding of phobic mechanisms. To expand on these advancements, we present data on the heterogeneity of neural patterns in spider phobia combined with various psychological dimensions of spider phobia, using spider-relevant stimuli of various intensities. Specifically, we have created a database in which 49 spider-fearful individuals viewed 225 spider-relevant images in the fMRI scanner and performed behavioral avoidance tasks before and after the fMRI scan. For each participant, the database consists of the neuroimaging part, which includes an anatomical scan, 5 passive-viewing and 2 resting-state functional runs in both raw and pre-processed form along with associated quality control reports. Additionally, a behavioral section includes self-report questionnaires and avoidance tasks collected in pre- and post-sessions. The dataset is well suited for investigating neural mechanisms of phobias, brain-behavior correlations, and also contributes to the existing phobic neuroimaging datasets with spider-fearful samples.

|  |  |
| --- | --- |
| Measurements | Anatomical and functional brain measurements, Online and in-person behavioral measurements. |
| Technology type(s) | Magnetic resonance imaging |
| Sample Characteristic-Organism | Homo sapiens |

## Background & Summary

Specific phobias are the most common anxiety disorders and can cause high levels of distress in affected individuals (e.g., Marks, 1987; Wardenaar et al., 2017). Spider phobia, which is defined by intense and exaggerated fear of spiders (American Psychiatric Association, 2013), is observed in a particularly large proportion of the population, with prevalence estimates ranging from 2.7% up to 9.5% (Fredrikson et al., 1996; Oosterink et al., 2009; Zsido, 2017). With the help of neuroimaging, researchers can gain insights into the neural underpinnings of such specific phobias. Particularly functional magnetic resonance imaging (fMRI) has revealed brain correlates of emotions and psychological dimensions relevant to spider phobia (for a review, see Hinze et al., 2021). Neuroimaging studies have identified a so-called “fear network” in the processing of spider-related stimuli, which consists of the amygdala, the insula and the anterior cingulate cortex (Hinze et al., 2021; Holzschneider & Mulert, 2011). Other studies have suggested that disgust, which has been linked to the prevention of contamination (e.g., Davey, 1994; Gerdes et al., 2009; Olatunji & McKay, 2006), is associated with overlapping but distinct neural activation compared to fear (Stark et al., 2007). As negative emotions, both fear and disgust are linked to avoidance behavior (e.g., Solarz, 1960), which is a key symptom of anxiety disorders (American Psychiatric Association, 2013). While phobic individuals tend to avoid the respective aversive stimulus, those who want to overcome their aversion may choose to approach the stimulus to expose themselves, resulting in an approach-avoidance conflict (Lewin, 1935). A variety of brain correlates of the approach-avoidance conflict have been proposed and include the inferior frontal gyrus, as well as the right dorsolateral prefrontal cortex (for details see Zorowitz et al., 2019). These findings can serve as a basis for further investigation of the specific underlying neural patterns in spider phobic individuals (Rosenbaum et al., 2020). Importantly, spider phobia is not a uniform condition (Knopf & Pössel, 2009). Individuals with spider phobia may have different levels of fear, disgust, avoidance behavior, and physiological responses (e.g., Schienle et al., 2005). This heterogeneity can make it challenging to identify consistent neural patterns across individuals. One goal of the current research is to address this heterogeneity issue by investigating psychological dimensions relevant to spider phobia and their brain correlates.

Besides neuroimaging, it is important to investigate spider phobia at the behavioral level. First, self-report questionnaires have been shown to indicate different levels of fear of spiders (e.g., Rinck et al., 2002), as well as related emotions and psychological constructs such as disgust, or state- and trait-anxiety (e.g., Laux et al., 1981; Schienle et al., 2002, 2010). Second, behavioral avoidance tests (BATs), in which spider-fearful individuals are asked to approach a spider, are important measures that can provide further insight into avoidance behavior (e.g., Antony et al., 2002; Muris et al., 1998). More recently, some computerized versions of BATs have been developed (e.g., Grill et al., 2023; Mühlberger et al., 2008). Despite the availability of these measures, knowledge of how behavioral data relate to brain data is limited. This knowledge could greatly benefit clinical applications and the development of tailored interventions (McNally, 2007).

Here, we present behavioral and MRI data of 49 individuals with fear of spiders. Specifically, we collected behavioral data consisting of a variety of self-report questionnaires that are highly relevant to fear of spiders, a physical behavioral avoidance test with a real spider, and a novel computerized behavioral avoidance test. Moreover, participants underwent an extensive MRI scan with a passive-viewing task, in which they were presented with images depicting spiders and neutral images. A resting-state scan was performed both before and after the passive-viewing task. Anatomical data were also acquired. After the MRI appointment, participants rated the spider images they had been presented with on a 101-point scale, according to “fear”, “disgust” and “willingness to approach”. Finally, participants underwent two follow-up assessments to examine a potential reduction in fear of spiders. As a result, the present study not only provides a large neuroimaging dataset, but also combines a broad variety of measurements, including both conventional and novel approaches. A subset of the data was analysized (but not released) in a previous study (Lor et al., 2023). Specifically, we used resting-state data from 37 out of 49 participants to compare functional connectivity before and after the passive-viewing task. We found decreased thalamo-cortical and increased intra-thalamic connectivity, suggesting that resting-state measures can be affected by prior emotion-inducing tasks and need to be carefully considered when detecting clinical biomarkers. The remainder of the data has not been previously used and the dataset is suitable for many questions investigating the neural mechanisms of phobias, fear processing, correlation between brain activity and phobic behavior as well as appraisal, interventions, and treatments, etc. This dataset also contributes to the growing collection of open MRI datasets, in particular it contains neural responses from spider-fearful individuals that are relatively difficult to access.

## Methods

### Participants

Fifty-one healthy, German-speaking individuals (41 female, 10 male) between the ages of 18 and 37 (mean = 22.65 ± 3.54) with self-reported fear of spiders and the willingness to overcome their fear participated in the study. Due to technical reasons, two participants had to be excluded from the dataset and the analysis, resulting in a total of 49 participants (40 female, 9 male; mean age = 22.55 ± 3.52, min = 18, max = 37). Participants were recruited via the university’s participant database, where they received an invitation and indicated their interest in participating. After signing up, they received an MRI safety questionnaire to evaluate their suitability to undergo an MRI examination and filled out the German Spider Fear Screening Questionnaire (SAS, Rinck et al., 2002; mean = 15.9 ± 4.4) where they had to score a minimum of 8 out of 24 points, indicating at least a moderate fear of spiders. Other exclusion criteria were self-reported past or current diagnosed psychiatric illnesses, pregnancy, or a history of alcohol or drug abuse. Participants received a financial compensation of 50€, with some receiving additional 5€ to compensate them for a pre-participation COVID-19 testing when required by the guidelines at the time. Data were collected between September 2021 and January 2023. The study was conducted in accordance with the Declaration of Helsinki and approved by the Ethics Committee of the University of Vienna (IRB numbers: 00584, 00657).

### Stimuli

The stimulus set used in the present study consisted of 225 images depicting spiders or spider-related content, and was a subset of the 313 images from a spider image database previously established by our lab (https://osf.io/vmuza/?view_only=93ff65b6629e4ae08174accfc6f3273b). These images were rated according to the dimensions of “fear”, “disgust”, and “willingness to approach” by individuals with self-reported fear of spiders in a previous study (Karner et al., 2024). To select 225 images from the 313 images that differed the most on these 3 psychological dimensions, ratings first underwent a whitening transformation to account for the different scaling of the ratings and their covariance using the “whitening” package in R (R Core Team, 2023). Then we used the “maximin” package to make a space-filling design using 225 images under the consideration of maximum-minimum distance. Some images depicted cobwebs, cartoon spiders or small spiders, while other images showed large spiders such as tarantulas or huntsman spiders, spiders eating prey, spiders in contact with human skin, or similar. All images had a size of 800×600 pixels and were upscaled to 1000×750 pixels for the passive-viewing fMRI scan.

### Experimental Procedures

The experiment consisted of a set of online surveys with self-report questionnaires, a behavioral avoidance test (BAT) with an actual spider, a computerized BAT, a passive-viewing MRI scan, ratings of spider images, as well as two follow-up sessions. An overview of the experimental design is provided in Figure 1.

**Figure 1.**
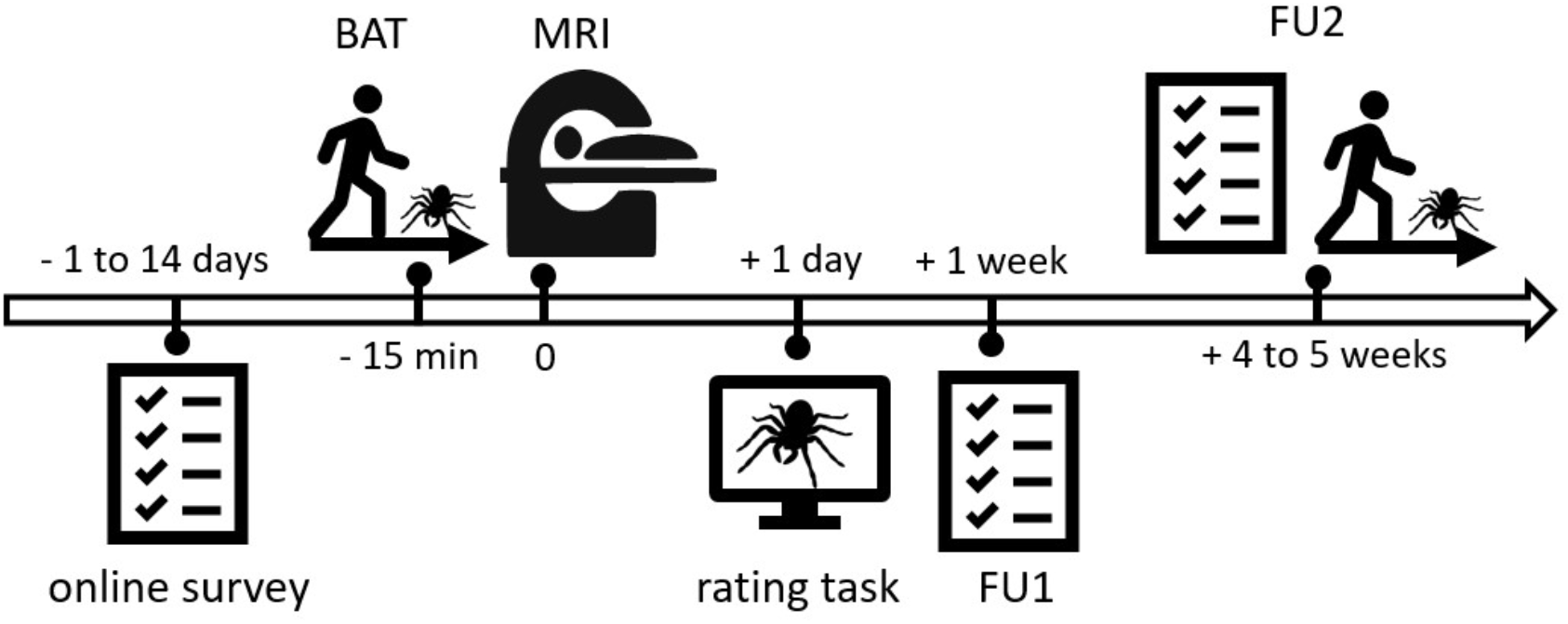
Overview of the experimental design. After confirming eligibility for participation in the study, participants received a link to an online survey with self-report questionnaires, that was filled out between 1 and 14 days prior to the MRI appointment. On the day of the MRI appointment, participants underwent a behavioral avoidance test (BAT) with a spider, and a computerized BAT. Approximately 15 minutes after completing the BATs, they underwent an MRI scan, during which they were presented with spider images on a screen. One day after the MRI appointment, participants received a link to an online image rating task, in which they rated the images they had been presented with according to “fear”, “disgust”, and “willingness to approach”. One week after the MRI appointment, participants engaged in a brief online follow-up (FU1), there they filled out a self-report questionnaire. Between 4 and 5 weeks after the MRI appointment, participants underwent a second follow up (FU2), which consisted of a self-report questionnaire, a BAT, and a computerized BAT.

### Self-Report Questionnaires

At a time point between one and 14 days prior to the MRI appointment, participants received a link to an online questionnaire on the SoSci platform of University of Vienna (Leiner, 2019), where they answered self-report questionnaires indicating their level of fear of spiders, specifically the German validated versions of the Fear of Spiders Questionnaire (FSQ; Szymanski & O’Donohue, 1995) and the Spider Phobia Questionnaire (Watts & Sharrock, 1984), that where introduced by Rinck et al. (2022), together with the German Spider Screening (Rinck et al., 2002). Additonally, participants filled out questionnaires indicating state- and trait anxiety (STAI; Laux et al., 1981), disgust propensity (FEE; Schienle et al., 2002) and disgust sensitivity (SEE; Schienle et al., 2010). Demographic information was also collected.

### Behavioral Avoidance Test

On the day of the MRI appointment, participants came to the University of Vienna, where they underwent a BAT, in which they were asked to approach a terrarium containing a spider. The spider was a real huntsman spider (*Heteropoda ssp*.; Latreille, 1804), which was inanimate and prepared in a natural-looking position. Participants were unaware that the spider was not alive. Before entering the room with the terrarium, they received the following instructions from the experimenter: *“In this room, we have a terrarium containing a spider. We would like to ask you to approach the terrarium up until a point where you still feel comfortable, and then stop and let us know. If you want to go all the way to the terrarium and feel comfortable, you may open the transparent lid, but you don’t have to do this*.*”* (translated from German) Then, the experimenter opened the door to the room with the terrarium and asked the participants to position their right foot behind a cross on the floor, which was 4.5 meters away from the terrarium. The experimenter remained standing in the door frame. While participants approached the terrarium, the distance covered was measured in centimetres using a laser distance meter that was always positioned in the same predefined spot and directed towards the participant’s leg. Specifically, we measured the start and end distance, to calculate the net distance that participants walked towards the terrarium. Two different terrariums (terrarium *A* and terrarium *B*) with identical spiders, but slightly different interior was used within the scope of the study to prevent potential biases due to static mental representations of the terrarium. Half of the participants approached terrarium *A* on the day of the MRI appointment and terrarium *B* on the day of the follow-up appointment, and vice versa.

### Computerized Behavioral Avoidance Test

In addition to the BAT, participants underwent a novel computerized BAT that was designed based on the work of (Peirce et al., 2019) and implemented in PsychoPy. In this task, participants needed to sharpen three highly blurred images of spiders up until a point where they still felt comfortable looking at them by dragging a slider on the screen. Although the scale was linear, with values from 1 to 101 stored, the sharpening of the images occurred in a logarithmic manner. The experiment was set up in PsychoPy and adapted from the script of Peirce et al. (2019). At the start of the computerized BAT, participants read the following instructions on the screen: *“On the following pages, you will see blurred, unrecognizable images of spiders. Use the slider to sharpen each image to the point where you still feel comfortable looking at it. Important: Keep the left mouse button pressed while moving the slider. As soon as you release the left mouse button, your answer will be saved. We will start with a practice round with a neutral image. This will be followed by the spider images. If you want to start, press the Space key*.*”* (translated from German) The practice image consisted of a chair on a wooden floor, followed by three spider images displaying large and hairy spiders. Contrary to the BAT, the experimenter left the room while participants engaged in the computerized BAT. During 11 computerized BATs (6 before the MRI appointment, 5 at follow-up 2) the experimenter remained present in the room with the participant. The order of performing the BAT first or the computerized BAT first was counterbalanced.

### Passive-Viewing MRI Scan

After completing the BAT and computerized BAT, participants underwent a brain scan at the University of Vienna’s MR Center. To reduce head motion, easily removable, non-constrictive piece of tape was placed on the participant’s forehead to provide tactile feedback as described by (Krause et al., 2019). Stimuli were presented to participants on an MR-compatible LCD screen (BOLDscreen 32 LCD for fMRI, Cambridge Research Systems) that was positioned at the head of the scanner bore. Participants were able to view the screen through a mirror mounted onto the head coil. First, a resting-state scan was performed during which participants were instructed to relax and keep their eyes open and directed at a fixation point. Subsequently, participants engaged in a passive-viewing task that consisted of five runs. In each run, participants were presented with 45 spider images, which displayed spiders and spider-related content, as well as with 15 neutral images depicting inanimate objects. Each image presentation lasted for 4 seconds and was followed by a presentation of the fixation point for 2-3 seconds. All images had a size of 1000×750 pixels. In each run, participants also underwent 6 catch trials, in which they were instructed to press a specific button on a button box that was positioned in their hand (e.g. *“Please press the left button now*.*”*). Once a catch trial was completed, the next image was presented to the participant. As an additional attention check, the experimenters used the live video of an eye tracker to see whether participants were keeping their eyes open, which was communicated to the participants beforehand. After runs 1, 3 and 5, participants were asked about their current levels of agitation and exhaustion, which they stated via the intercom on a scale from 0 to 10. The five passive-viewing runs were followed by another resting-state scan which was identical to the first resting-state scan. This was followed by a structural scan, which marked the end of the scanning session. The MRI data therefore consist of 7 functional runs, 5 passive viewing runs and two resting state runs, as well as a structural scan.

### Image Ratings

One day after the MRI appointment, participants received an email with links to the University of Vienna’s SoSci (Leiner, 2019) platform, where they were asked to rate all spider images that had been presented to them during the passive viewing fMRI scan. The ratings were administered in three blocks (A, B. and C), each consisting of 75 images. The images were presented in their original size of 800x600 pixels. Participants were allowed to take breaks between the blocks, with the requirement to complete all three blocks within 48 hours after receiving the links. However, ratings that were administered within one week after the fMRI appointment were also accepted by the research team. The image rating procedure was as follows: Each block started with a practice round, followed by the actual image ratings. In each image rating trial, an image was presented on the screen for 3 seconds. Then the image disappeared, and 3 questions and corresponding rating scales were presented on the screen.

(1) “How much fear does this picture elicit in you?”; (2) “How much disgust does this picture elicit in you?”; (3) “How close could you come to the content shown in the picture, if you wanted to overcome your aversion?” (translated from German). Ratings were made by moving a continuously adjustable slider on a 101-point visual analog scale. In each trial, participants were given 12 seconds to administer the three ratings. To proceed with the next image, they clicked “Next” in the bottom-right corner on the screen. Each block contained a break questionnaire, in which participants stated their current levels of fear, physical arousal, disgust, boredom and exhaustion in subjective units of distress (Wolpe, 1969) on a scale from 0-10. Additionally, each break questionnaire contained a bogus item to test the participants’ attention (e.g., Meade & Craig, 2012).

### Follow-up Assessments

One week after the MRI appointment, participants received a link to an online questionnaire where they were asked to fill out the FSQ a second time (follow-up 1). A total of 38 out of 49 participants completed follow-up 1. Follow-up 2 took place between 4 and 5 weeks after the MRI appointment. Participants returned to the University of Vienna, where they first filled out the FSQ a third time and then underwent the BAT and computerized BAT for a second time. Thirteen participants filled out the FSQ after having completed the BAT and computerized BAT. Participants were then asked to fill out a brief feedback sheet asking them about their thoughts on the study, and whether they noticed anything special. A total of 46 out of 49 participants completed follow-up 2. Out of all the participants, only one person expressed doubt about the authenticity of the spider. The feedback sheet marked the end of the participation in the experiment.

### MRI Acquisition

All functional and structural scans were collected using a 3T Siemens MAGNETOM Skyra MRI scanner (Siemens, Erlangen, Germany) with a 32-channel head coil, located at the University of Vienna. Functional data were acquired with an interleaved mode, using a T2*-weighted echo planar imaging sequence with multiband acceleration factor 4, TR = 1250 ms, TE = 36 ms, voxel size = 2 mm * 2mm * 2mm, flip angle = 65 degrees, field of view (FOV) = 192 mm * 192 mm * 145 mm, 56 slices without slice gap to cover the full brain, and anterior-posterior phase encoding. The total scan duration for each resting state run was 9:00 mins, and 7:20 mins for each passive viewing run. After the functional runs, a field map was collected with a short echo time of 4.92 ms and a long echo time of 7.38 ms.

Structural data for each participant was acquired using a single-shot, high-resolution MPRAGE sequence with an acceleration factor of 2 using GRAPPA with TR = 2300 ms, TE = 2.43 ms, voxel size = 0.8 mm * 0.8 mm * 0.8 mm, flip angle = 8 degrees, and field of view (FOV) = 263 mm * 350 mm * 350 mm. The total duration of the structural data acquisition was 6:35 mins.

### MRI Preprocessing

The pre-processed data included in this dataset was performed with fMRIPrep 20.2.6 (Esteban et al., 2019), which is based on Nipype 1.7.0 (Gorgolewski et al., 2011).

### Anatomical data preprocessing

The T1-weighted (T1w) images were first corrected for intensity non-uniformity (INU) using N4BiasFieldCorrection (Tustison et al., 2010), and used as T1w-reference throughout the workflow. Then, the T1w-reference was skull-stripped with antsBrainExtraction.sh workflow, using OASIS30ANTs as the target template. Brain tissue segmentation of cerebrospinal fluid (CSF), white-matter (WM) and gray-matter (GM) was performed on the brain-extracted T1w using the software fast (FSL 5.0.9, RRID:SCR_002823, Zhang et al., 2001). Volume-based spatial normalization to the standard space (MNI152NLin2009cAsym) was performed through nonlinear registration using antsRegistration (ANTs 2.3.3), with brain-extracted versions of both T1w reference and T1w template.

### Functional data preprocessing

For each of the 7 functional runs per subject, the following procedures were performed: First, a reference volume and its skull-stripped version were generated using a custom methodology of fMRIPrep. A B_0_-nonuniformity map (i.e. fieldmap) was estimated based on a phase-difference map calculated with a dual-echo GRE (gradient-recall echo) sequence, processed with a custom workflow of SDCFlows. The field map was then co-registered to the target EPI (echo-planar imaging) reference run and converted to a displacement field map with FSL’s fugue and other SDCflows tools. Based on the estimated susceptibility distortion, a corrected EPI reference was calculated for a more accurate co-registration with the anatomical reference. The reference was then co-registered to the T1w reference using flirt, and the co-registration was configured with nine degrees of freedom to account for distortions remaining in the reference. Head-motion parameters (transformation matrices, and six corresponding rotation and translation parameters) were estimated before any spatiotemporal filtering using mcflirt (Jenkinson et al., 2002). BOLD runs were slice-time corrected to 0.571s using 3dTshift (Cox & Hyde, 1997). The fMRI time-series were resampled onto their original, native space by applying a single, composite transform to correct for head-motion and susceptibility distortions. Also, the fMRI time-series were resampled into standard space, generating a preprocessed functional run in the MNI152NLin2009cAsym space. Several confounding time-series were calculated based on the preprocessed BOLD: framewise displacement (FD), DVARS and three region-wise global signals. FD and DVARS were calculated for each functional run, both using their implementations in Nipype, and the three global signals were extracted within the CSF, the WM and the whole-brain masks. Additionally, a set of physiological regressors were extracted to allow for component-based noise correction (CompCor, Behzadi et al., 2007). Principal components were estimated after high-pass filtering the preprocessed fMRI time-series for the two CompCor variants: temporal (tCompCor) and anatomical (aCompCor). tCompCor components were then calculated from the top 2% variable voxels within the brain mask. For aCompCor, three probabilistic masks (CSF, WM and combined CSF+WM) were generated in anatomical space. From these masks, a mask of voxels likely to contain a volume fraction of GM was subtracted and then resampled in BOLD space. The head-motion estimates calculated in the correction step were also placed within the corresponding confounds files. All resamplings were performed with a single interpolation step by composing all the pertinent transformations. (i.e. head-motion transform matrices, susceptibility distortion correction when available, and co-registrations to anatomical and output spaces). Gridded (volumetric) resamplings were performed using antsApplyTransforms (ANTs), configured with Lanczos interpolation to minimize the smoothing effects of other kernels (Lanczos, 1964). Non-gridded (surface) resamplings were performed using mri_vol2surf (FreeSurfer).

Many internal operations of fMRIPrep use Nilearn 0.6.2 (Abraham et al., 2014; RRID:SCR_001362), mostly within the functional processing workflow. For more details of the pipeline, see the section corresponding to workflows in fMRIPrep’s documentation (https://fmriprep.org/en/20.2.6/workflows.html).

## Data Records

The MRI dataset is organized according to the Brain Imaging Data Structure (BIDS; Gorgolewski et al., 2016) specification (version 1.0.1) and evaluated with the MRI Quality Control tool (MRIQC) version 1.4.0 (Esteban et al., 2017). All imaging data, including preprocessed data with fMRIprep version 20.2.6 (under the “derivatives/fmriprep” directory) and quality assessment reports with MRIQC version 1.4.0 (under the “derivative/mriqc” directory) are available on OpenNeuro (https://openneuro.org/datasets/ds004630). The behavioural data, the corresponding codebooks and the computerized BAT are available on OSF (https://osf.io/w4s2h/). We set up a GitHub repository (https://github.com/univiemops/spider20-fmri-data) to share the data descriptions and the preprocessing and analysis scripts.

## Technical Validation

### Quality Assessment

To assess the quality of the current dataset, we used an open-source BIDS application, MRIQC, to automatically generate Image Quality Metrics (IQMs) and visual quality reports for individual and group levels. Individual functional reports were generated per subject per run to show the mosaic view of the average BOLD signal map, standard deviation from average BOLD signal map, summary plot of motion-related indices, and IQMs. The IQMs include commonly used measures of spatial and temporal signal quality (e.g., signal-to-noise ratio (SNR), temporal SNR (tSNR, Figure 2), DVARS) and measures of artifacts (e.g., framewise displacement (FD, Figure 3), number of dummy scans). The individual anatomical reports include a mosaic zoomed-in brain mask map, a background noise map, IQMs including noise measurements. The group functional and anatomical reports visualize the IQMs across participants by displaying a box plot per measure. These reports graphically show the range and outliers of quality measures and can be used to quickly compare participants. All reports and accompanying JSON and TSV files are shared on OpenNeuro.

**Figure 2.**
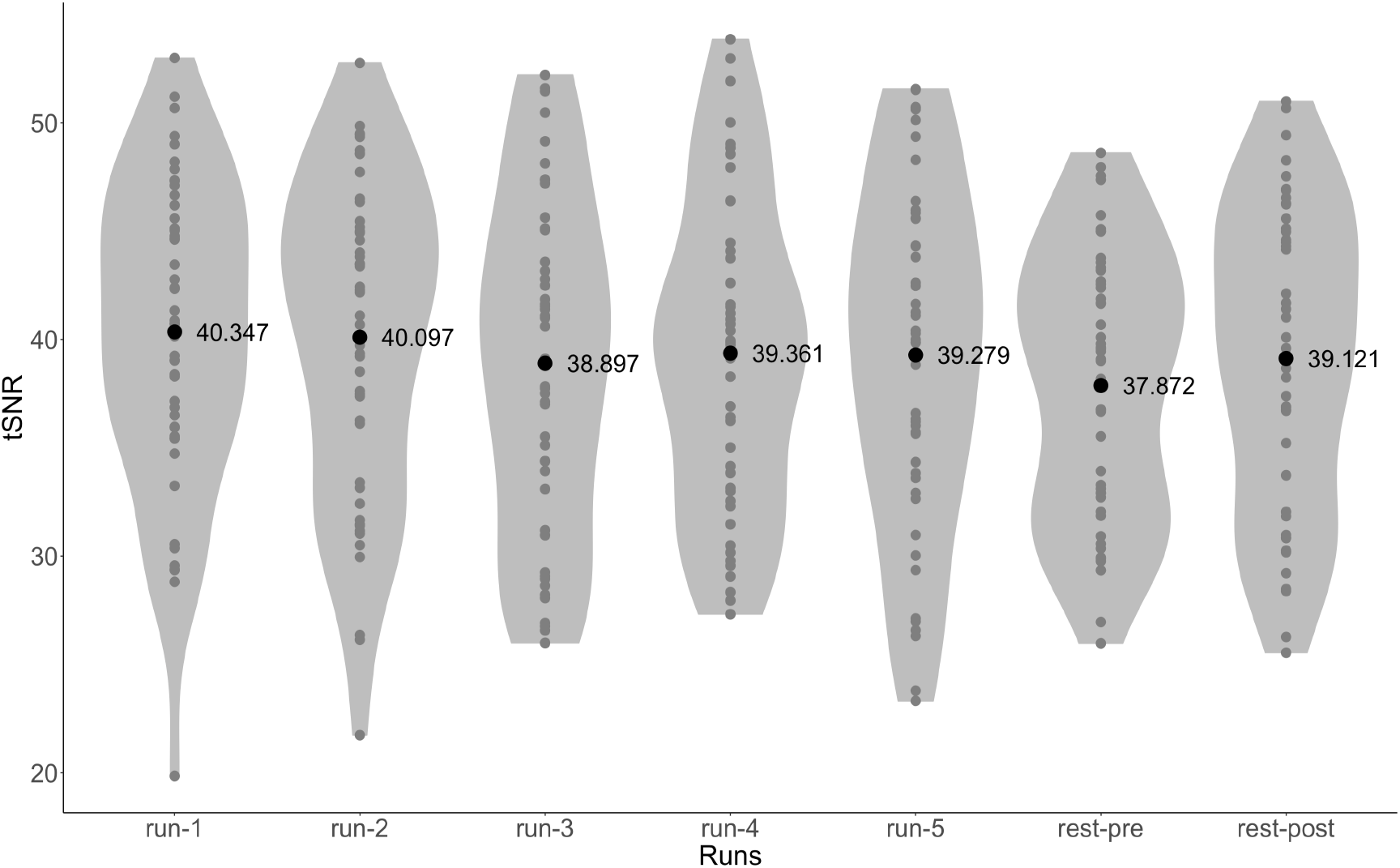
Temporal SNR of each run across all participants. Gray dots represent the individual tSNR for that run. Black dots and associated values are the mean tSNR across all participants for that run. The mean whole-brain tSNR across participants for all runs was 39.28 ± 7.20.

**Figure 3.**
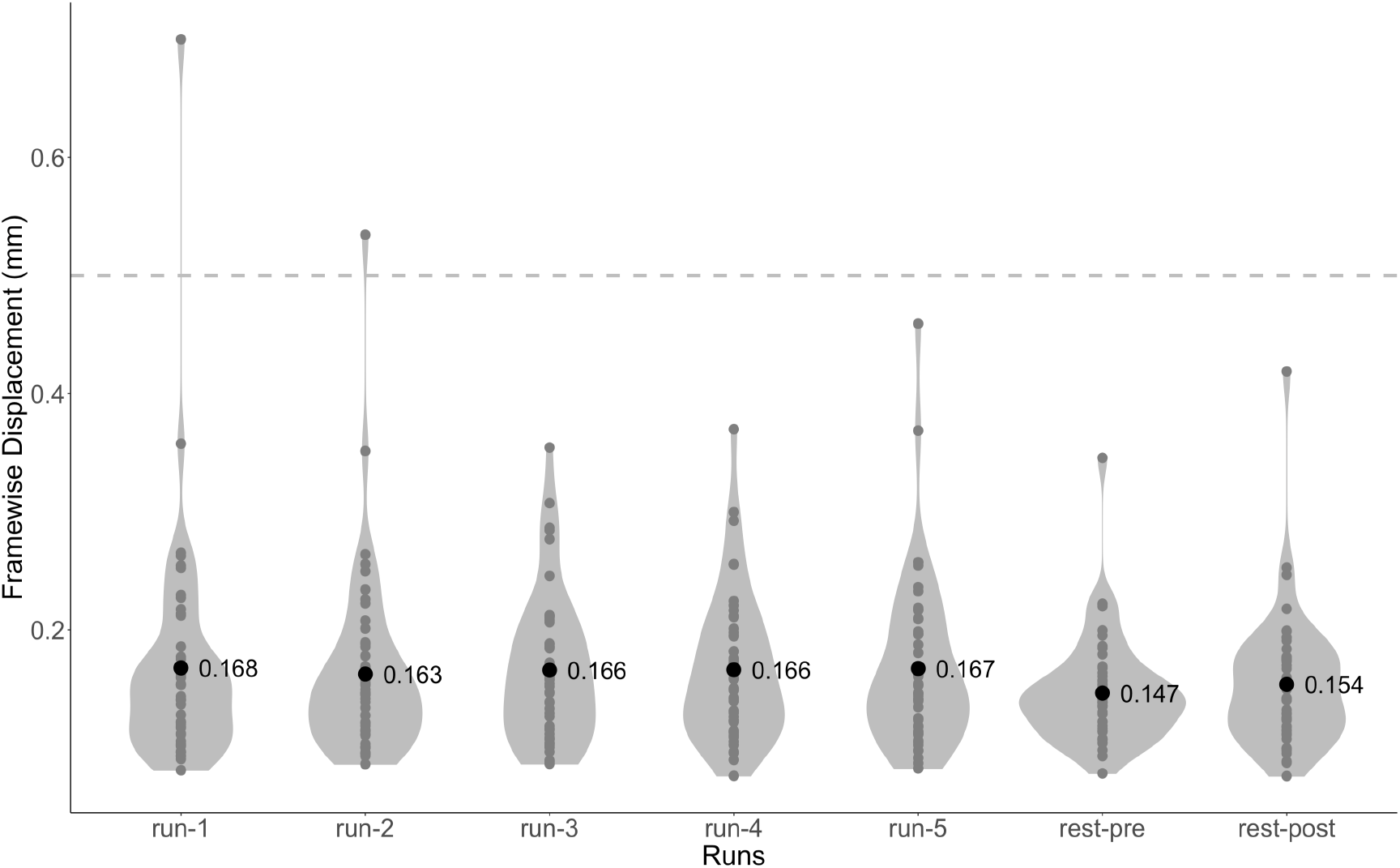
Framewise displacement of each run across all participants. Gray dots represent the individual FD for that run. Black dots and associated values are the mean FD across all participants for that run. Participant movement is low in this dataset, only 2 runs of one participant had higher than 0.5mm FD. The mean FD across participants for all runs was 0.16 ± 0.07 mm.

**Figure 4.**
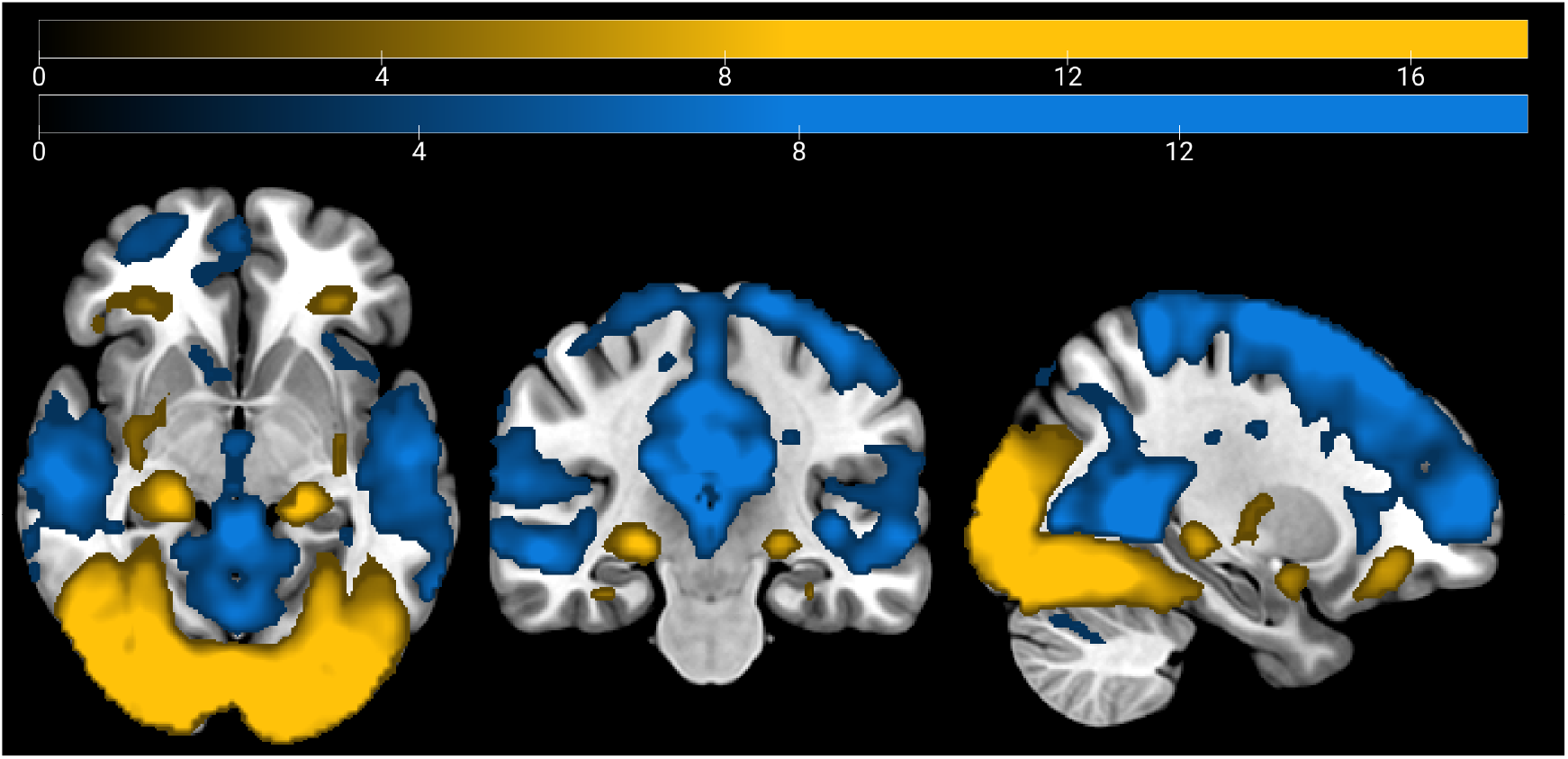
Brain activation clusters for stimulus presentation vs. implicit baseline (p<0.001 for peak threshold and p<0.05 for FWE-cluster correction). Blue: negative activation; yellow: positive activation. N=49.

### Basic analysis of stimuli vs. implicit baseline

To further validate the dataset, we performed a basic general linear model (GLM) analysis to show brain activation during stimulus presentation compared to an implicit baseline. This analysis should find positive voxel activation in visual areas and task-related regions, such as the hippocampus and the amygdala. For this purpose, we used the fMRIprep-preprocessed passive-viewing functional scans and Statistical Parametric Mapping (SPM12; Wellcome Trust Centre for Neuroimaging, London, United Kingdom) for the GLM analysis. Runs for which the maximum framewise displacement (FD) exceeded 5mm were excluded from the analysis (2 runs of one participant). For the first-level analysis, we specified two regressors, one for the onsets of stimulus presentation (including spider images and neutral images) and one for the onsets of the button-press catch trials. Each stimulus was modelled as a boxcar function, with lengths matching each stimulus duration, and convolved with the hemodynamic response function. The six motion realignment parameters were added as nuisance regressors. We performed a one-sample t-test second-level analysis, one for positive activation and one for negative activation, using the first-level activation maps that correspond to the “stimuli” regressor from the 49 participants. We applied a family-wise error corrected cluster threshold of p < 0.05, with height threshold set at p<0.001. As expected, we found positive activation in visual areas, as well as fear-related regions such as the bilateral amygdala, hippocampus and putamen. Conversely, we found negative activation in regions associated with resting-state such as parts of the default-mode network among other regions. This outcome is consistent with the previous literature on similar experimental paradigms (Botvinik-Nezer et al., 2019).

## Code Availability

All code is available in the GitHub repository (https://github.com/univiemops/spider20-fmri-data). The code includes scripts for experiment presentation (executed in Python and largely relied on PsychoPy (Peirce, 2007)), data sorting, preprocessing and quality control (executed in shell scripting), and analyses (executed in Matlab).

## Acknowledgements

FM was funded by the Austrian Science Fund (FWF) [10.55776/ESP133]. KK was funded by the Austrian Science Fund (FWF) [10.55776/ESP286]. FS was supported by the Swiss National Science Foundation (BSSG10_155915, 100014_178841, 32003B_166566), the Foundation for Research in Science and the Humanities at the University of Zurich (STWF-17-012), and the Baugarten Stiftung.

## Author contributions

MZ: Conceptualization, Formal Analysis, Investigation, Data Curation, Validation, Visualization, Writing - Original Draft. AK: Conceptualization, Methodology, Software, Formal Analysis, Investigation, Data Curation, Writing - Original Draft. KK: Conceptualization, Data Curation, Writing - Review & Editing. SS: Investigation, Writing - Review & Editing. DS: Conceptualization, Writing - Review & Editing. RS: Methodology, Resources, Writing - Review & Editing. FM: Software, Writing - Review & Editing. CL: Conceptualization, Methodology, Software, Validation, Formal Analysis, Investigation, Data Curation, Writing - Original Draft. FS: Conceptualization, Project Administration, Methodology, Supervision, Funding Acquisition, Resources, Writing - Review & Editing.

